# MHC Attention: Identifying HLA-E presented cancer antigens through deep learning and high-throughput screening

**DOI:** 10.64898/2026.06.08.730987

**Authors:** Ethan Fast, Manjima Dhar, Gunsagar S. Gulati, Millicent Ku, Binbin Chen

**Affiliations:** Vcreate Inc., Menlo Park, CA, USA; Department of Medical Oncology, Dana-Farber Cancer Institute, Boston, MA, USA

**Author notes:** These authors contributed equally to this work (Ethan Fast, Manjima Dhar).

## Abstract

HLA-E presented cancer peptides can be promising cancer therapy targets, as HLA-E is minimally polymorphic and widely expressed across human populations and cancer types. However, systematic discovery of cancer associated HLA-E peptides has been constrained by sparse training data and the technical difficulty of HLA-E immunopeptidomics. Here we develop an integrated HLA-E antigen discovery platform combining a deep learning prediction model, pooled mammalian cell screening, peptide-HLA-E stability validation, and mass spectrometry. We introduce MHC Attention, a neural network that learns allele-level attention over candidate MHC alleles in multi-allele immunopeptidomics datasets, enabling direct training on patient-derived MHC peptide data. Screening an approximately 6,000-peptide HLA-E library identified stable HLA-E-presented peptides and generated HLA-E-specific training data that improved prediction performance of MHC Attention. Combining our screening assays and improved prediction algorithm, we discovered novel HLA-E-presented cancer peptides, including candidates derived from ETV4, WT1, RNF43 and BMP8A, with orthogonal support from stability assays or immunopeptidomics. These results establish a scalable framework for HLA-E peptide target discovery and provide candidate targets for broadly applicable peptide–HLA-directed cancer immunotherapies. MHC Attention 2.0 can be accessed online via https://vcreate.io/mhcattention.

## Introduction

Immunotherapies against peptide-major histocompatibility complexes (pMHCs) offer a strategy to drug intracellular targets. However, the field has largely focused on classical MHC class I molecules, HLA-A, -B, and -C, which are highly polymorphic and therefore restrict treatment to allele-defined patient populations (*3–6*). Both FDA-approved pMHC-targeting drugs, tebentafusp-tebn and afamitresgene autoleucel, are restricted to patients with HLA-A*02 subtypes (20-50% of the US, depending on the ethnic sub-population) (*7–9*). In contrast, the non-classical class I molecule HLA-E has remained comparatively underexplored despite its minimal polymorphism: HLA-E\**01:01 and HLA-E\**01:03 account for more than 99% of HLA-E alleles worldwide (*10*). Immunotherapies that drug peptide-HLA-E targets could enable global clinical trials with broader patient populations (*11, 12*).

HLA-E is also compelling due to its direct involvement in tumor immune evasion. In its canonical role, HLA-E presents peptide fragments derived from leader sequences of HLA class I, such as VMAPRTVLL (VL9 peptides) (*13*). These peptides engage CD94/NKG2A receptors to restrain natural killer (NK) cells and subsets of CD8+ cytotoxic T cells (*14*), and many cancer types upregulate HLA-E to reinforce this inhibitory axis against NK cell attacks (*15–19*). Beyond this canonical role, emerging evidence indicates that HLA-E can also present cancer-associated peptides distinct from VL9 peptides, creating opportunities for therapeutic targeting. For example, recent work has shown that vaccination can elicit HLA-E-restricted CD8+ T cell response against tumor-associated peptides including prostatic acid phosphatase (PAP), Wilms’ tumor-1 (WT1), and mesothelin, providing a proof of principle that HLA-E-restricted antitumor immunity can be therapeutically exploited (*20*).

Unfortunately, it remains difficult to discover HLA-E-presented ligands across genes of clinical interest (*21*). Immunopeptidomics, which has transformed antigen identification for classical HLA-I, is poorly suited to HLA-E: surface expression of HLA-E is substantially lower than that of classical class I molecules, the peptidome is dominated by canonical VL9 peptides, and few HLA-E-specific antibodies are available for immunoprecipitation (*14, 22, 23*). Biochemical binding and stability assays can interrogate defined peptide sets, but HLA-E complexes with non-canonical peptides are often weakly stable, making readouts noisy (*24, 25*). Computational prediction can in principle bridge this throughput gap, however, the performance of existing MHC prediction algorithms depends on large, high-quality training sets that do not yet exist for HLA-E, where validated ligands remain sparse and presentation rules diverge from those of classical alleles (*1, 26–29*).

To address these challenges, we developed a new platform for HLA-E target discovery that combines machine learning guided peptide nomination with screening based on cell libraries. Candidate peptides are selected from genes of interest using MHC Attention, a novel deep neural network that learns an attention mechanism to deconvolute the likely presenting MHC allele in patient mass spectrometry data. Selected peptides are then screened in a high-throughput mammalian cell library based on single-chain trimers composed of a peptide, β2-microglobulin, and HLA-E. In this assay format, peptides that fail to stabilize HLA-E produce unstable complexes with low surface expression, converting peptide-dependent HLA-E stability into a scalable cellular readout. Data from this cell library readout can be used to select peptides for validation in MHC stability assays and to finetune MHC Attention, improving the algorithm’s ability to differentiate HLA-E-presented peptides (*30, 31*).

We applied this platform to peptides from 141 cancer-associated genes identified from transcriptomic data, known IEDB-reported HLA-E presenters (*32*), and predicted peptides from the human proteome. We then validated the efficacy of the discovery platform with conventional HLA-E stability assays, confirming the discovery of many previously unrecognized HLA-E-presented peptides. Finally, MHC Attention was retrained on these data and the updated model was used to screen and re-rank peptides from the same set of cancer-associated genes. We tested the resulting top-ranked candidates and discovered additional novel HLA-E presenters that may be suitable for future cancer immunotherapy development.

## Results

### An integrated discovery and validation platform for HLA-E-presented cancer antigens

To identify cancer antigens presented by HLA-E, we established an integrated platform that combines transcriptomic prioritization, peptide presentation modeling, pooled functional screening, and orthogonal MHC stability assays (**Fig. 1A**). We combined both literature reports and RNA-sequencing data from normal and malignant tissues to nominate cancer-selective protein-coding genes, then enumerated candidate peptides from these proteins for presentation scoring. To rank this search space, we developed a novel MHC class I prediction algorithm capable of modeling peptide presentability across MHC alleles (MHC Attention, **Fig. 2**) and applied it to peptides derived from clinically relevant cancer-associated proteins (**Fig. 1B**). Because HLA-E presentation is strongly constrained by peptide-dependent complex stability, we next built a mammalian cell-based high-throughput assay in which candidate peptides are expressed in peptide–β2-microglobulin–HLA-E single-chain trimers, such that stabilizing peptides promote higher surface HLA-E expression whereas poorly presented peptides yield unstable complexes with lower surface signal (**Fig. 3**).

**Figure 1.**
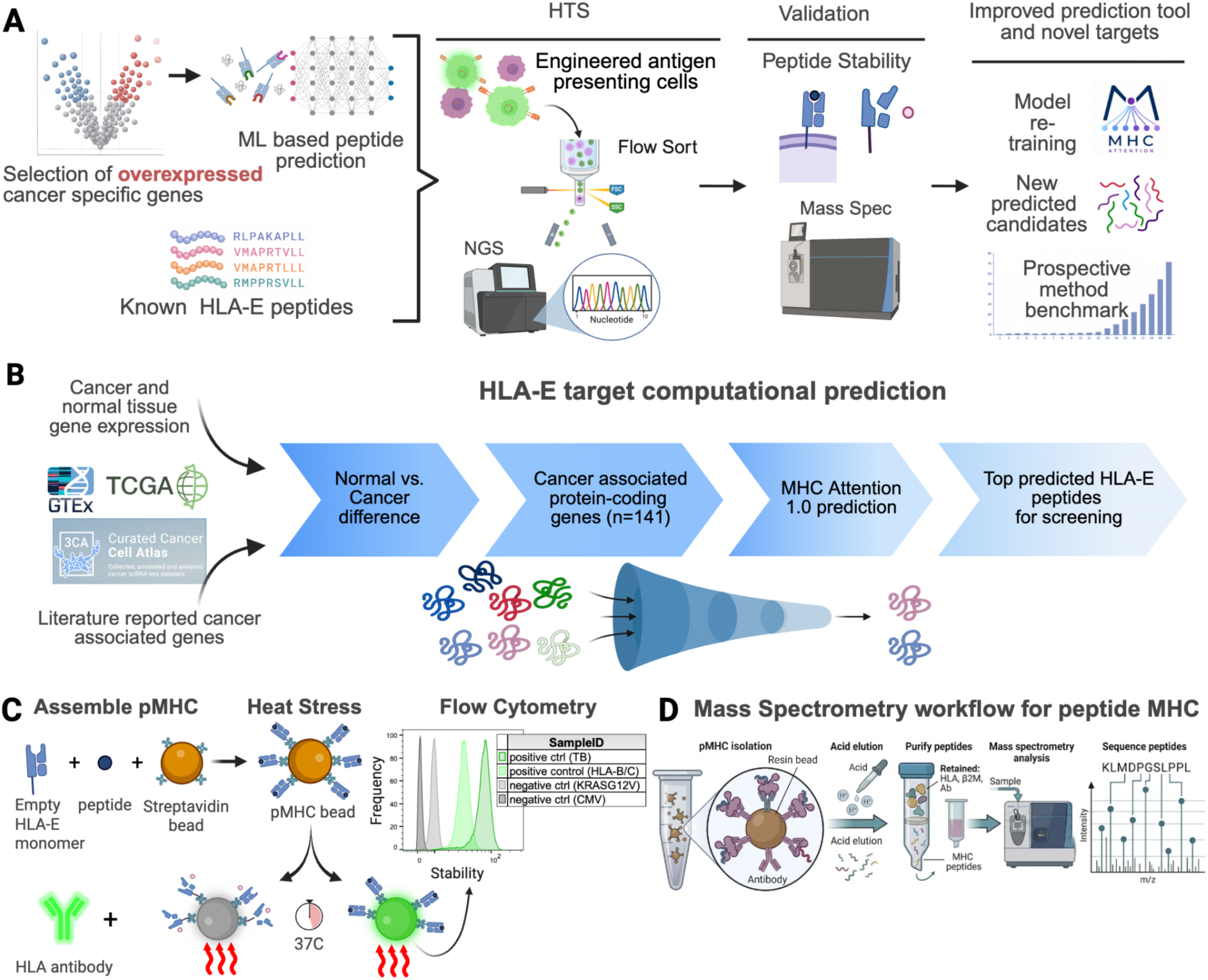
Integrated workflow for HLA-E target discovery and validation. **A.** Schematic of the overall discovery platform. Cancer-selective genes were determined using bulk and single-cell RNA-sequencing data from both normal and cancer tissues. Cancer-associated protein-encoding genes were then screened with computational presentation prediction models for potential HLA-E presentable peptides. We next applied high-throughput screening with engineered antigen-presenting cells, flow sorting, and NGS-based analysis to identify stable HLA-E peptide candidates. Candidate hits were validated by peptide stability assays and HLA-E immunopeptidomics by mass spectrometry. **B.** Computational workflow for HLA-E target selection. The Cancer Genome Atlas Program (TCGA), the Genotype-Tissue Expression (GTEx), the Curated Cancer Cell Atlas (3CA), and Cancer Antigenic Peptide (CAPED) databases were used to derive 141 cancer-associated genes differentially expressed between cancer and normal tissues. Peptide 9mers from these genes were scanned using MHC Attention 1.0, and high scoring peptides were selected for inclusion in a high-throughput cell-based HLA-E screening assay. **C.** Bead-based HLA-E stability assay. Peptide-loaded biotinylated HLA-E monomers were assembled on streptavidin coated beads, subjected to heat stress at 37° C for 60 minutes, and stained with anti-HLA antibody for flow cytometric measurement of residual stable pMHC complexes. Clear separation of flow signals was observed between positive and negative control peptides. TB-RLPAKAPLL and VL9-VMAPRTVLL were used as positive controls, and KRAS-VVVGAVGVGK and CMV-NLVPMVATV as negative controls. **D.** HLA-E immunopeptidomics workflow. HLA-E complexes from cancer cells were isolated by immunoprecipitation. Bound peptides were then acid-eluted and purified, and peptide identities were determined by LC–MS/MS.

**Figure 2.**
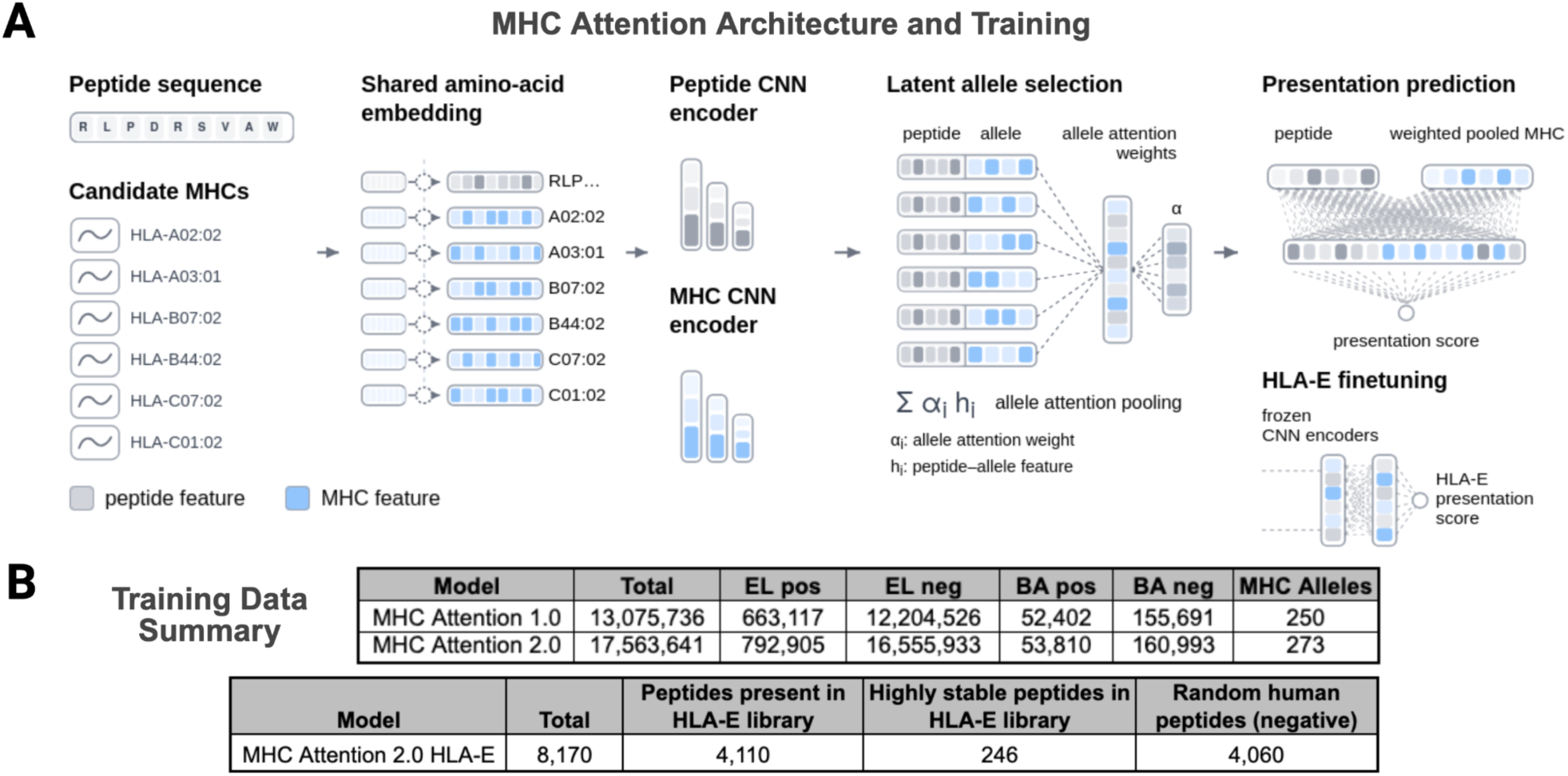
Overview of the MHC Attention peptide presentation prediction model. **A.** Architecture of MHC Attention. MHC Attention is a deep neural network that predicts MHC class I presentation. Its central design feature is MHC allele attention, where weighting over candidate alleles is learned during training on multi-allele examples as a form of pointwise attention. Conventional mass spectrometry data has six MHC alleles and no label for the true presenter, so this architecture enables training directly on such data. MHC Attention encodes peptides and alleles through independent convolutional neural network (CNN) encoders with a shared amino acid embedding, learns an allele attention mechanism over candidate alleles, and produces a presentation score between 0 and 1 after processing the attended candidate set through a final dense layer. MHC Attention was also finetuned on HLA-E data from a cell library (Fig. 3) with frozen encoders and a dedicated HLA-E presentation head. **B.** MHC Attention training datasets. Three training datasets were used to train model versions of MHC Attention. MHC Attention 1.0 was trained on 13,075,736 peptides across 250 MHC alleles, consisting of eluted ligand (EL) and binding (BA) data. MHC Attention 2.0 was trained on an expanded dataset of 17,563,641 peptides across 273 alleles. MHC Attention 2.0 HLA-E was trained as a new prediction head on top of frozen MHC Attention 2.0 encoders using continuous HLA-E stability values for 4,110 peptides from an HLA-E cell library as well as 4,060 additional random human peptides assigned negative labels.

**Figure 3.**
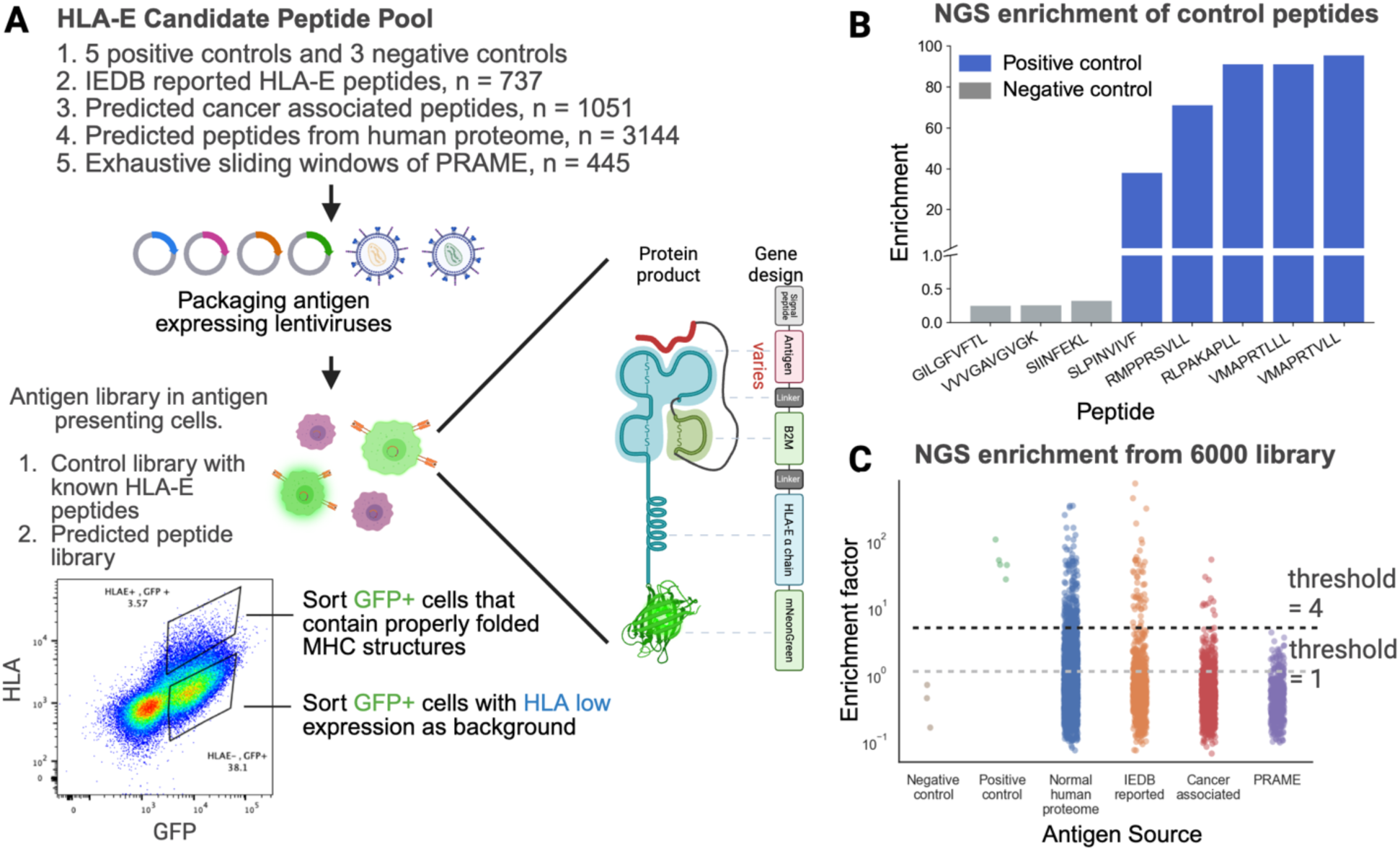
High-throughput cell-based screening assay for HLA-E peptide stability. **A.** Design of the pooled HLA-E screening assay. The peptide library was composed of five peptide sources: control peptides (5 positives and 3 negatives), IEDB-reported HLA-E peptides (n = 736), peptides from cancer-associated genes/proteins (n = 1051), predicted peptides from the normal human proteome (n = 3144), and PRAME-derived peptides (n = 437). Library DNA was packaged into lentiviral vectors encoding peptide-B2M-HLA single-chain trimer constructs. These constructs encode a single protein consisting of a signal peptide, antigen peptide, β2-microglobulin, GGGGS linker, HLA-E alpha chain (Y84A), and C-terminus green fluorescent protein (GFP). HLA-null K562 cells transduced with the construct lentivirus become GFP-positive. Constructs with poor HLA-E presenting peptides are unstable and lead to low surface HLA levels for K562 cells. Populations with GFP+ high surface HLA level and GFP+ low HLA level were sorted separately and sequenced for the composition of the peptides. HLA-E stability for each peptide can be estimated by the ratio of peptide abundance in HLA high vs. low cell populations (enrichment). **B.** Sequencing-based enrichment analysis of control peptides. Following sorting, known positive control peptides (VL9-VMAPRTVLL, VL9-VMAPRTLLL, TB-RLPAKAPLL, INTS1-RMPPRSVLL, ORF1AB-SLPINVIVF) showed preferential enrichment in the HLA-high fraction relative to negative control peptides (M1-GILGFVFTL, KRAS-VVVGAVGVGK, OVA-SIINFEKL), supporting the ability of the assay to recover good HLA-E presenting peptides. **C.** Distribution of enrichment values. Stability enrichment analysis was performed per peptide source across the HLA-E 6,000 peptide library. Enrichment was calculated as the ratio of a peptide’s fractional representation in the HLA-high sorted population relative to its fractional representation in the HLA-low sorted population, after sorting and sequencing. An enrichment threshold of 4 was used to select candidate peptides for follow-up validation. 244/3144 peptides with enrichment >4 were observed for predicted normal human proteome peptides, 61/737 for IEDB-reported peptides, 15/1051 for predicted cancer-associated peptides, 1/445 for PRAME-derived peptides, and 5/5 for positive controls.

We used this platform to screen a library of ∼6,000 peptides comprising both candidates from cancer-associated proteins and literature-reported HLA-E controls, followed by fluorescence-based sorting and sequencing-based deconvolution to quantify peptide enrichment in HLA-high versus HLA-low populations (Assay **Fig. 3**; MHC Attention **Fig. 4**). Candidate hits were then validated using two approaches: a bead-based peptide stability assay that directly measures persistence of peptide-loaded HLA-E complexes after heat challenge (**Fig. 1C, 5**), and HLA-E immunopeptidomics by immunoprecipitation and LC–MS/MS to confirm endogenous peptide recovery from cancer cells (**Fig. 1D**). We then used the cell-based screening results to expand HLA-E-specific training data and finetune the prediction model with additional training (**Fig. 2, 4**). Finally, we conducted a new round of HLA-E target nomination and validation across cancer associated genes with the improved computational model. We compared our fine-tuned HLA-E prediction algorithm against the state-of-the-art tool by predicting and validating novel HLA-E peptides (**Fig. 6**).

**Figure 4.**
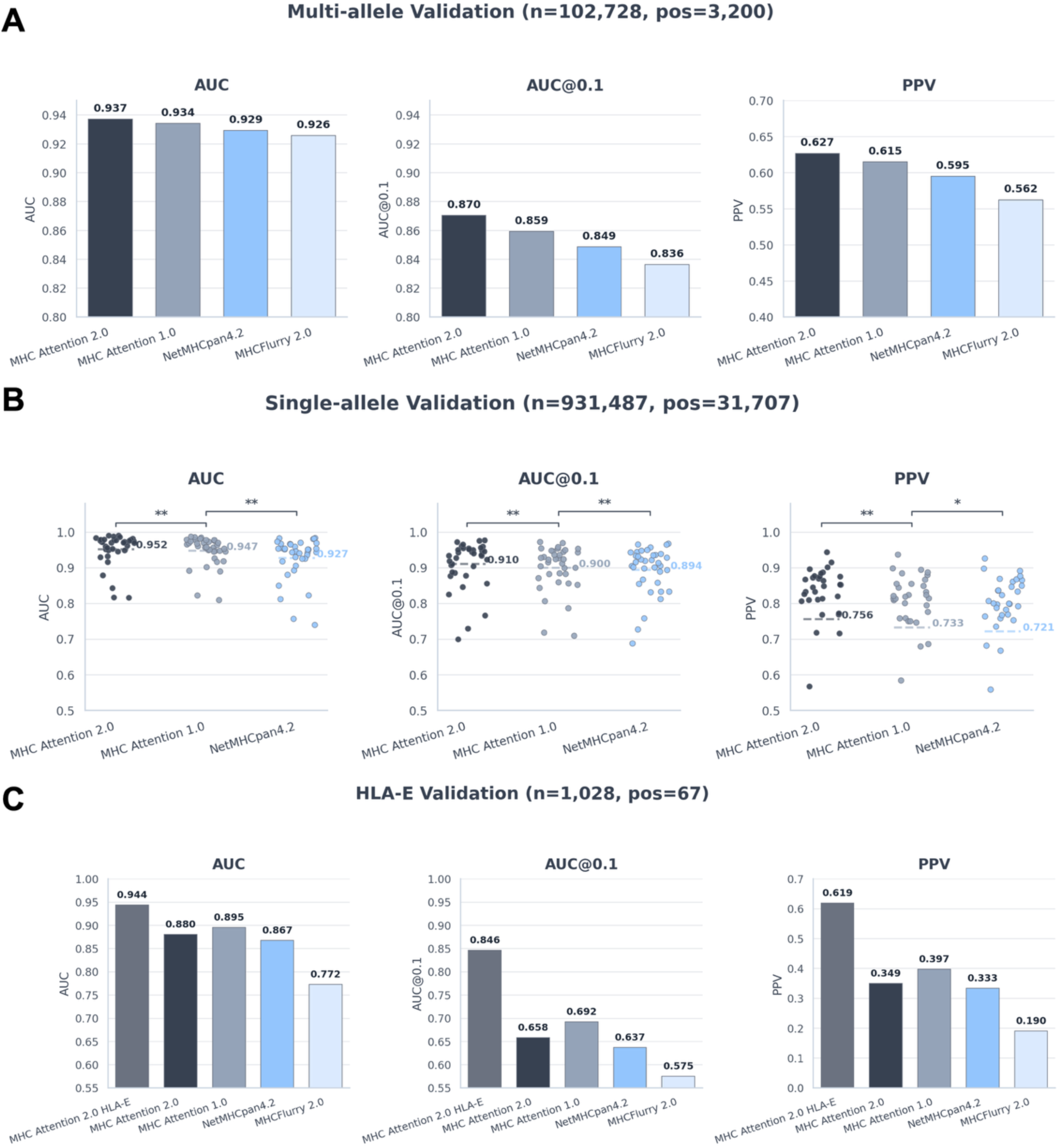
Benchmark performance of MHC Attention 2.0 and HLA-E-specific model refinement. We compared several versions of MHC Attention with NetMHCpan4.2 and MHCflurry 2.0. MHC Attention 2.0 HLA-E was trained on additional HLA-E library presentation data generated in this study. Metrics shown are area under the receiver operating characteristic curve (AUC), partial AUC at false-positive rate 0.1 (AUC0.1), and positive predictive value (PPV), which we report as precision within the top *k* predictions where *k* = 0.95 × (number of positives in the dataset). All comparisons use unnormalized prediction scores. **A.** Performance on the filtered multi-allele validation benchmark assembled from the Sarkizova et al. dataset (*1*) after removal of training overlap with all methods. Across 102,728 peptides with 3,200 positive peptides, MHC Attention 2.0 achieved the strongest performance across all evaluation metrics. **B.** Performance on the single-allele validation benchmark composed of 931,487 peptides and 31,707 positives across 36 alleles (*2*). Each point represents one allele. MHC Attention 2.0 showed the strongest performance across all metrics. MHCflurry 2.0 was excluded from this benchmark due to near-complete overlap with its training data. P-values were computed with a paired Wilcoxon signed-rank test across alleles (*, p < 0.05; **, p < 0.01). Detailed allele level performance break-down was included in **Supplementary Table 7**. **C.** Performance on the HLA-E validation benchmark derived from held-out HLA-E cell-based screening data. Positives were defined as peptides with HLA-E enrichment above 4, which gave 67 positives among 1,028 peptides. MHC Attention 2.0 HLA-E outperformed all other methods.

### Identification of cancer associated protein-coding genes

To define a tractable search space for HLA-E target discovery, we assembled cancer-associated protein-coding genes from two complementary tracks: a curated track seeded from a known cancer-antigen database, and a discovery track based on bulk and single-cell transcriptomic analysis. Throughout, we restricted analysis to protein-coding genes, excluding pseudogenes and non-coding transcripts (**Fig. 1B**).

For the discovery track, we integrated three transcriptomic resources. GTEx was used to establish normal-tissue baseline expression (*33*), taking the median TPM across donors for each tissue, with particular emphasis on critical organs relevant to on-target toxicity (full tissue list in Methods). TCGA bulk RNA-seq data (*34*) was used to quantify tumor expression, again as the median TPM across samples within each cancer type, and to prioritize indications with high unmet clinical need (full cancer-type list in **Methods**). To add cell-type resolution beyond bulk comparisons, we incorporated the Curated Cancer Cell Atlas (3CA), a harmonized collection of cancer single-cell RNA-seq datasets (*35*). Within 3CA, we computed a malignancy-discrimination score for each gene as the area under the receiver operating characteristic curve (AUC) distinguishing malignant from non-malignant cells, including endothelial, fibroblast and immune populations in the tumor microenvironment, averaged across datasets (see **Methods** for details).

Genes in the discovery track were prioritized using three filters: greater than fourfold higher median expression in tumor than in normal tissue, a malignancy-discrimination AUC above 0.6, and median expression below 5 TPM in critical normal organs including brain, heart, lung, kidney, and liver. This transcriptome-based screen identified a set of cancer-associated genes with potentially favorable therapeutic windows (*33–35*).

For the curated track, we drew on established cancer-antigen resources, primarily the Cancer Antigenic Peptide Database (CAPED) (*36*), whose genes have documented support for producing cancer-associated MHC peptides. We then applied translational and mechanistic considerations across both tracks. Because direct cell-surface modalities such as antibodies, antibody-drug conjugates, and conventional bispecifics are already well-suited to membrane proteins, we de-prioritized surface antigens such as LY6G6D and CDH6 in favor of intracellular proteins, where peptide-HLA targeting offers a clearer comparative advantage, and we gave higher priority to genes with existing clinical or translational precedent.

Combining the curated and discovery tracks yielded a final set of 141 cancer-associated genes (**Supplementary Table 1**). These included antigens such as PRAME and MAGEA4, which have already supported clinical TCR-based cell therapy programs, as well as targets such as MIF, which has substantial therapeutic rationale in oncology despite being less advanced as a peptide-HLA target (*37–40*).

### A deep learning model to better predict peptide MHC Class I presentation

The 141 cancer-associated genes we selected contain tens of thousands of candidate peptides, so further prioritization is required before library screening. Deep learning models can predict peptide presentation well for classical MHC class I alleles, and efficiently score large numbers of peptides, but their performance for HLA-E remains less well established.

To address this gap, we developed MHC Attention 1.0, a neural predictor of MHC class I presentation that learns jointly from single-allele and multi-allele immunopeptidomics data. Its primary innovation over existing MHC presentation prediction models is the treatment of the presenting allele in multi-allele samples as a latent variable, with attention over candidate alleles learned explicitly during training. This is important because most mass spectrometry datasets contain six patient alleles and no label for the true presenter; MHC attention can train end-to-end on such data and learn to deconvolute the likely presenting allele. A peptide and its candidate alleles are encoded in the network though convolutional neural network (CNN) encoders with a shared amino acid embedding (*41*). These representations pass through an allele attention mechanism, creating a pooled MHC representation from which presentation is predicted. (**Fig. 2A**). We hypothesized that this end-to-end architecture for allele deconvolution over multi-allele samples (e.g., all primary cancer tissues) might improve MHC presentation prediction performance overall, and in particular on predictions for HLA-E.

We trained MHC Attention 1.0 on 13,075,736 examples of MHC peptide presentation (715,519 positive, 12,360,217 negative) across 250 alleles curated by Reynisson et al (**Fig. 2B**) (*26*). Each example consisted of a single peptide, one or more MHC alleles, and a presentation label. We then evaluated the model on two general peptide presentation benchmarks, comparing it with two leading methods, NetMHCPan4.2 and MHCflurry 2.0 (*2, 27*). The first benchmark evaluates area under the receiver operating characteristic curve (AUC), AUC up to 0.1 false positive rate (AUC0.1), and positive predictive value (PPV) over held-out multi-allele samples from primary cancer samples (*1*). All methods gave strong performance on this benchmark, however MHC Attention 1.0 gave the strongest performance in terms of overall AUC, AUC0.1, and PPV (**Fig. 4A**). MHC Attention 1.0 demonstrated strongest differentiation in PPV (0.615 vs 0.595 for NetMHCPan4.2 and 0.562 for MHC Flurry 2.0). The second benchmark similarly evaluated the same metrics over 931,487 peptides in Sarkizova et al. single-allele data (**Fig. 4B**). For this benchmark we did not compare with MHCflurry 2.0 due to >90% overlap in its training data, but MHC Attention 1.0 again performed stronger than NetMHCPan4.2 over AUC (+0.0207, p=6.0e-9), AUC0.1 (+0.00581, p=0.0051), and PPV (+0.0115, p=0.039).

These results suggest that MHC Attention 1.0 offers stronger performance than existing models, though by a relatively small margin. Performance differences show up more clearly in stricter metrics like PPV and AUC0.1. These metrics, however, are quite valuable for candidate peptide nomination, where high positive predictive value (PPV) is desirable.

### Developing cell based high-throughput screening for HLA-E presentation

HLA-E is understudied relative to other MHC alleles, and the number of known presenting peptides is small and includes many peptides with limited evidence. To enhance the field’s understanding of HLA-E presentation, we wanted to evaluate reported peptides as well as candidate HLA-E presenting peptides from our analysis of cancer-associated genes.

Evaluating thousands of peptides for HLA-E presentation requires a more scalable assay than conventional MHC peptide stability testing. To functionally test candidate peptides in a cellular context, we developed a pooled high-throughput screening platform based on peptide–β2-microglobulin–HLA-E single-chain trimer constructs expressed in engineered antigen-presenting cells (**Fig. 3A**). In this format, peptides that fail to support stable HLA-E complex formation yield reduced surface HLA expression, whereas stabilizing peptides promote high surface HLA-E levels (*42*). Each construct contains a signal sequence, antigen peptide, β2-microglobulin, HLA-E α chain, and a C-terminal mNeonGreen reporter (GFP) linked by flexible GGGGS linkers. Following previously reported MHC class I single-chain trimer design principles, we introduced the Y84A mutation to accommodate the peptide–linker junction at the F pocket while preserving peptide-dependent differences in MHC stability, unlike Y84C-based disulfide trapping approaches that can artificially stabilize weakly presented peptides (*14, 43*).

The assay works by sorting GFP-positive cells into surface HLA-high and surface HLA-low fractions and quantifying peptide representation in each population by targeted mRNA sequencing (**Fig. 3A**). Enrichment is defined as the ratio of a peptide’s fractional abundance in the HLA-high population relative to the HLA-low population.

Using this approach, we developed a library comprised of approximately 6,000 peptides from five sources: common control HLA-E peptides (5 positives and 3 negatives) (*13*), IEDB-reported HLA-E peptides (n=737), peptides derived from cancer-associated genes (n=1051, >98 percentile for MHC Attention 1.0), predicted peptides from the normal human proteome (n=3144, >99.9 percentile cutoff for MHC Attention 1.0), PRAME-derived peptides (n=445) (**Fig. 3A**).

We validated the assay by examining the enrichment level of positive and negative control peptides. All five well-known HLA-E peptides included in the assay achieved 20-103 fold enrichment in the high HLA surface level cell population (**Fig. 3B**, TB-RLPAKAPLL, VL9-VMAPRTVLL, VL9-VMAPRTVLL, INTS1-RMPPRSVLL, and ORF1AB-SLPINVIVF) (*13*). Further, all three negative control peptides we included (M1-GILGFVFTL, KRAS-VVVGAVGVGK, and OVA-SIINFEKL) demonstrated enrichment values below 1. This clear separation confirmed that the assay could distinguish peptides that stabilize HLA-E from peptides that do not.

### Screening and validation of reported and predicted HLA-E peptides

To identify HLA-E presenting peptides from our 6000 peptide library, we transduced 5 million HLA-null K562 cells with the library lentiviruses at a low Multiplicity of Infection (MOI<0.3). We sorted around 100,000 K562 cells from both HLA high and HLA low cells from GFP+ cells post HLA E library construction. Antigen peptide coding region mRNA from these two populations were amplified and sequenced with RT-PCR and bulk sequencing with Illumina MiSeq.

Only a fraction of IEDB reported HLA-E peptides (61/737, 8.3%) met a stringent 4x enrichment cutoff (**Fig 3C, Supplementary Table 2**), despite the strong enrichment of our positive controls (5/5). We found a similar ratio (244/3144, 7.7%) of predicted peptides enriched at that cutoff in the human proteome, and fewer in cancer associated genes (15/1051, 1.4%) due to the lower score cutoff that we deliberately adopted (98 percentile for cancer associated vs 99.9 percentile for the human proteome) to include a relatively large number of cancer associated peptides from a more restricted set of cancer associated genes.

We then used a conventional pMHC stability assay to validate selected candidates, including a set of likely cancer-associated peptides as well as other candidates from genes of particular interest, including, PRAME-derived peptides and several other novel HLA-E peptides nominated independently of cancer association. In this pMHC stability assay, individual peptides were loaded onto biotinylated, peptide-receptive HLA-E monomers (**Fig. 1C**). Then the resulting pMHC complexes were bound to streptavidin-coated beads, followed by a heat challenge at 37 °C to destabilize weakly bound complexes. Intact pMHC complexes were then quantified by flow cytometry using an anti-HLA antibody, such that higher post-heat signal reflected greater peptide-dependent stabilization of HLA-E. We validated this assay with positive and negative control peptides. The stability of a peptide was quantified by the proportion (%) of pMHC beads with HLA signal above background (see **Methods**).

Out of 30 peptide candidates we tested with conventional stability assays, we observed peptides with higher enrichment in the screening assay exhibited higher stability (**Supplementary Table 4**). The median stability of peptides with greater than 4 fold enrichment was 27.8%, compared with 0.57% for peptides with low enrichment (**Fig. 5B**, p=6e-3). This strong but imperfect agreement is likely due to the fact that the cell library screening assay is based on single-chain trimers, which can have minor physiologic differences from natural HLA-E. The most stable peptides were candidates derived from WT1, RNF43, and LGSN.

**Figure 5.**
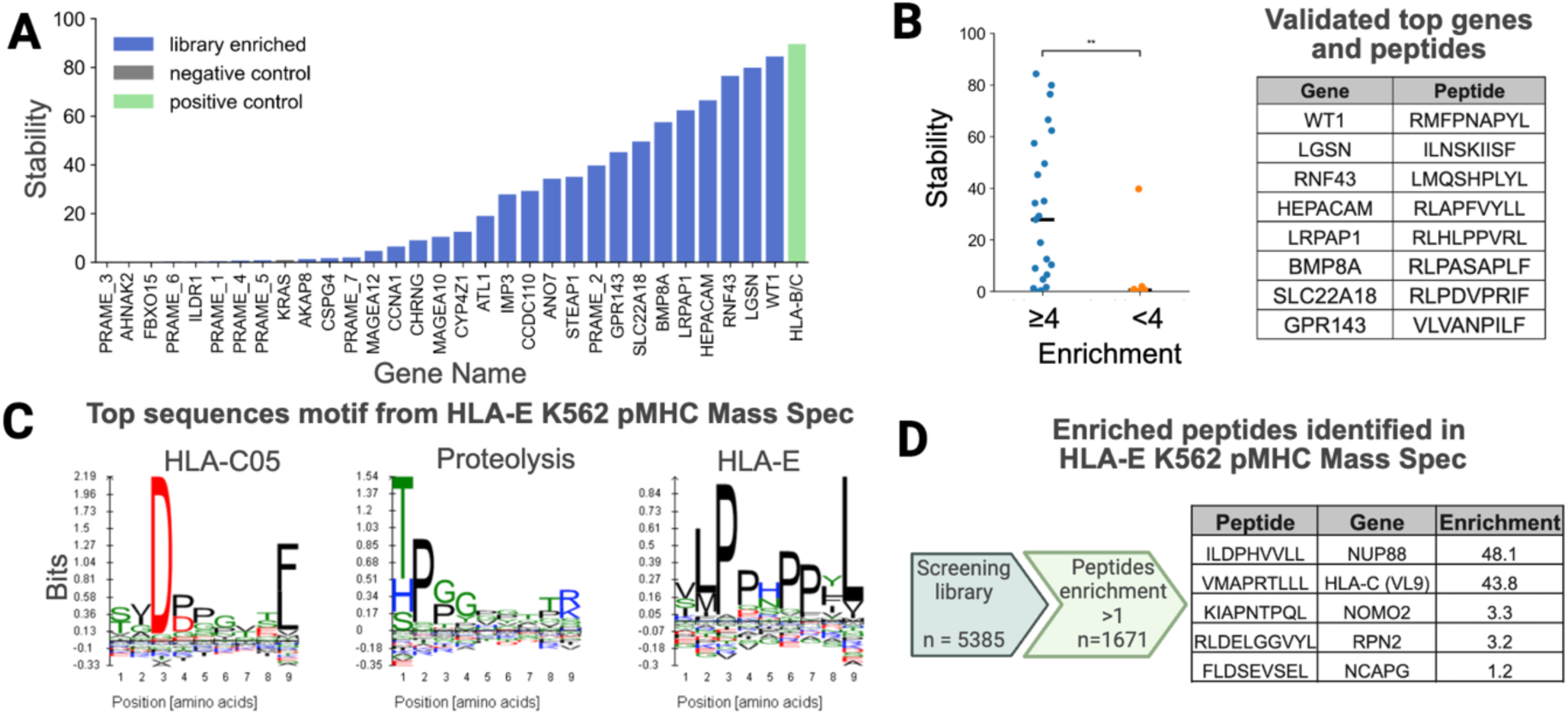
Experimental validation of HLA-E screening hits by peptide stability assay and mass spectrometry. **A.** Validation of candidate peptides by HLA-E stability assays. Candidate peptides identified from pooled screening were validated individually using peptide–HLA-E stability measurements. Peptides with stronger enrichment in the pooled screen generally showed higher HLA-E stability: 37% of the hits achieved greater than 30% flow-based stability score. **B.** Relationship between validation stability and cell screening enrichment. Stability measurements were examined for all peptides selected from the HLA-E library, broken down by peptides with high library enrichment (>4) and peptides with lower enrichment. These groups showed significantly different stability values, with a median stability among high enrichment peptides of 28% and a median stability among lower enrichment peptides of 0.57% (**, two-tailed two-sample unequal variance t-test, p = 6e-3). **C.** Sequence motif analysis from HLA-E immunopeptidomics. Peptide sequences (8-12mers) recovered from HLA-E K562 cells MS immunopeptidomics experiments were clustered using GibbsCluster. Motifs in three clusters matched known motifs for HLA-C*05, proteolysis products, and HLA-E. Representative 9mer sequence logo plots are shown. **D.** HLA peptides enriched in the cell-based screening that validated in HLA-E K562 immunopeptidomics. Out of 1671/5385 peptides with enrichment values greater than 1 in the cell screening library, five were detected in HLA-E K562 immunopeptidomics: Nup88-ILDPHVVLL, VL9-VMAPRTLLL, NOMO2-KIAPNTPQL, RPN2-RLDELGGVYL, NCAPG-FLDSEVSEL. K562 cells express 1745/3182 (54.8%) genes in the screening library at TPM greater than 1.

### Profile natural HLA-E peptide repertoire via MS

To further confirm our library screening approach, we investigated whether positive hits from the library screening can be detected by mass spectrometry based HLA-E immunopeptidomics. We first developed a K562 cell line with ultralow HLA Class I levels (HLA-null K562) and then introduced HLA-E transgenes via lenti-transduction (see **Methods** for details). These HLA-E K562 cells can be processed with conventional HLA Class I immunoprecipitation workflows (HLA Class I antibody clone W6/32) as most HLA molecules in these cells are HLA-E.

Working with Alithea Bio, we processed the HLA-E K562 cells and analyzed recovered peptides by LC–MS/MS. We identified 1,160 unique peptides (7-13 amino acids long) from the HLA-E K562 immunopeptidomics (**Supplementary Table 3**). We applied GibbsCluster (*44*) clustering and observed a motif for the main cluster of peptides consistent with HLA-E binding preference (**Fig. 5C**). Interestingly, we also observed a cluster of peptides exhibiting HLA-C05 sequence motifs. Given K562 cells carry HLA-C*05:01, this indicates some HLA-E K562 cells still expressed moderate levels of HLA-C. Out of 1671 peptides with enrichment values greater than 1 in the cell screening library, five were detected in HLA-E K562 immunopeptidomics (**Fig. 5D**) though not all genes in the cell screening library were expressed by K562 cells.

### Further improve MHC Attention HLA-E prediction performance by incorporating new data

We next investigated whether we could improve the performance of MHC Attention 1.0 using the new HLA-E associated peptide data. We trained a new version of the model, MHC Attention 2.0, on an expanded set of 17,563,641 examples across 273 alleles curated by Nilsson et al. (**Fig. 2B**) (*2*). We explored various ways of integrating the new HLA-E data with this expanded dataset. To preserve our ability to evaluate the model on unseen examples, we also held-out 20% of this new HLA-E data for testing.

We found that fine-tuning a base MHC Attention 2.0 model gave the best predictive performance on internal validation. In particular, we trained the classifier head with soft labels targeting enrichment scores on the HLA-E data from the screening library, leaving the existing peptide and MHC encoder and attention weights unchanged. We also incorporated additional random human peptides as negatives sampled evenly across peptide lengths (**Fig. 2**). These additional negatives approximately double the size of the fine-tuning data.

Overall, the HLA-E finetuned model gave stronger predictive performance than other models on the held-out HLA-E data across all metrics (**Fig. 4C**). The HLA-E finetuned model improved most dramatically on its ability to rank positives via PPV (+0.27 vs next highest) and AUC0.1 (+0.188 vs next highest). MHC Attention 1.0 also outperformed NetMHCPan4.2 and MHCflurry 2.0 on this benchmark, though score differences between models were much smaller.

On the more general MHC Class I benchmark set, we found that MHC Attention 2.0 consistently outperformed MHC Attention 1.0 across AUC, AUC0.1, and PPV by a small margin, in both the single-allele and multi-allele conditions (**Fig. 4A, 4B**). The strongest improvements were in PPV, with +0.012 PPV on multi-allele and +0.023 PPV in the single-allele condition.

### Benchmark HLA-E prediction performance on novel cancer peptide candidates

To further investigate whether the improvements in HLA-E specific training for MHC Attention 2.0 were helpful and identify additional cancer-specific HLA-E presenters, we performed a computational screening of all 9-11mer peptides (n=71,038) over a subset of the cancer associated genes we earlier selected (n=63, **Supplementary Table 1**). To better isolate the value of the HLA-E specific training over existing prediction algorithms, we performed this computational screening with both MHC Attention 2.0 HLA-E and NetMHCpan4.2, an existing leading algorithm in the field. We ran both algorithms over all peptides and selected the top 20 according to each algorithm’s presentation score for stability testing. There was no overlap in the peptides selected by MHC Attention and Netmhcpan4.2 in their top 20 ranking.

The stability assay identified 5 peptides with meaningful HLA-E presentation (**Fig. 6B, Supplementary Table 5**). The most stable presenter was an 11mer peptide from ETV4 identified by MHC Attention. We found 25% (5/20) precision for the peptides selected by MHC Attention and 0% (0/20) precision for the peptides selected by Netmhcpan4.2. Notably, the ETV4-derived peptide KLMDPGSLPPL and MATR3-derived peptide ILGPPPPSF were also supported by MS immunopeptidomics (**Fig. 6C, 6D**). Given the low sensitivity of MS immunopeptidomics, ETV4-KLMDPGSLPPL and MATR3-ILGPPPPSF are likely abundant in cancer HLA-E complexes.

**Figure 6.**
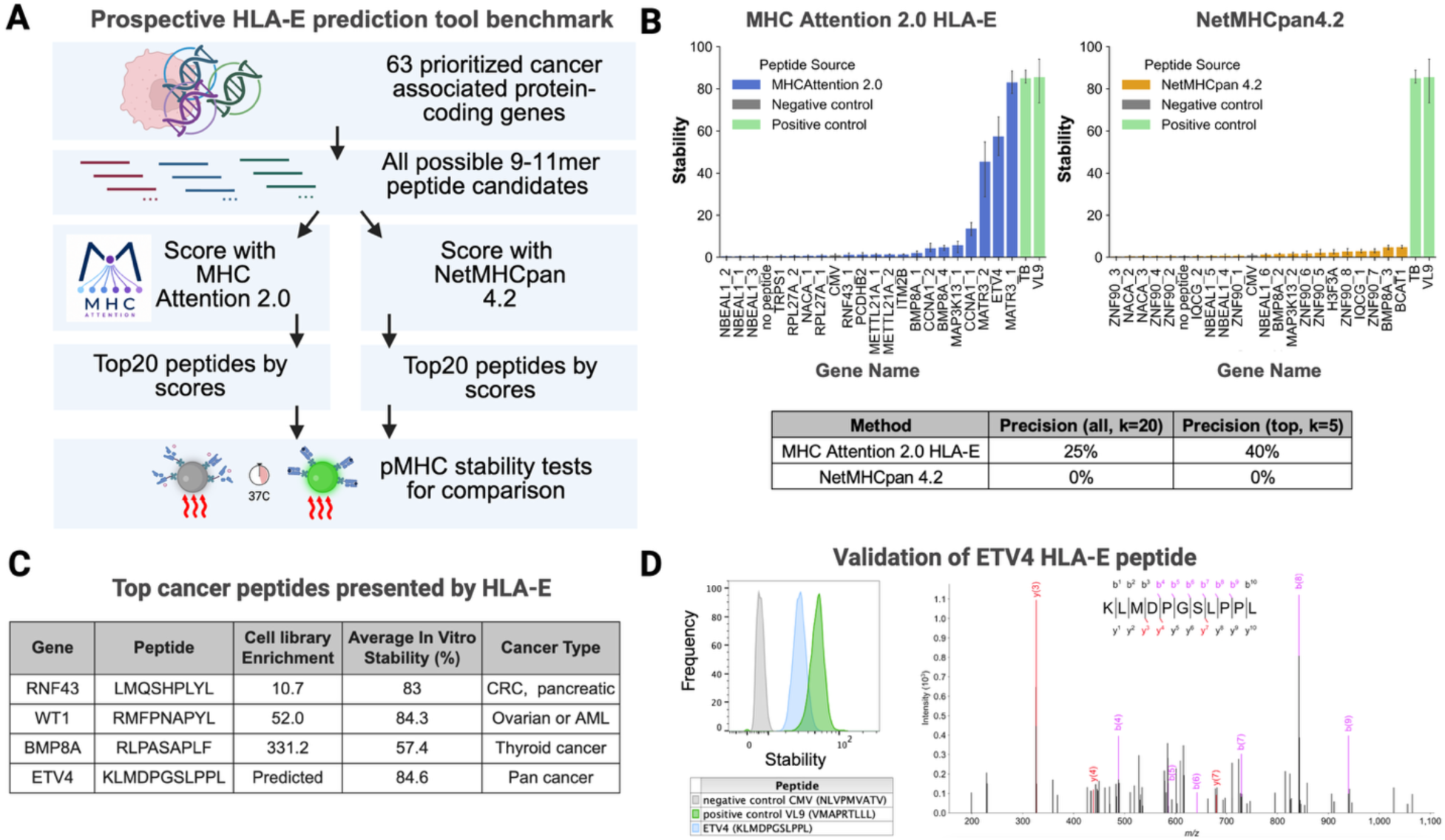
Benchmarking MHC Attention 2.0 HLA-E performance and identifying novel HLA-E presenting cancer-associated peptides. A. Evaluating MHC Attention 2.0 HLA-E and NetMHCpan4.2 for HLA-E peptide discovery. 63 cancer associated protein encoding genes were selected and 9-11mer sliding windows (n=71,038) were evaluated with computational models for HLA-E presentation. The twenty highest scoring peptides from MHC Attention 2.0 HLA-E and NetMHCpan4.2 were selected for stability validation. B. HLA-E stability results for the top twenty peptides selected by MHC Attention 2.0 HLA-E and NetMHCpan4.2. Five peptides demonstrated >5% HLA stability, all of which were selected by MHC Attention 2.0: MATR3-ILGPPPPSF, MATR3-ILGPPPPSFHL, CCNA1-SLMEPPAVLLL, MAP3K13-AMGNHPSPKL, ETV4-KLMDPGSLPPL. C. Top HLA-E-presented cancer peptides for therapeutic development. Peptides with positive HLA-E stability were further evaluated for differential cancer expression, strength of stability, and therapeutic potential. Four peptides were selected based on these criteria: RNF43-LMQSHPLYL, WT1-RMFPNAPYL, BMP8A-RLPASAPLF, and ETV4-KLMDPGSLPPL. In vitro stability values for RNF43, WT1 and ETV4 are reported as the mean of ≥3 independent experiments conducted over time. D. Experimental evidence for ETV4-KLMDPGSLPPL presentation by HLA-E. HLA-E stability for ETV4-KLMDPGSLPPL was measured by flow cytometry. The ETV4 peptide showed high stability (HLA level) after heat challenge compared to a negative control. The same ETV4 peptide was independently discovered in the HLA-E K562 cell MHC immunopeptidomics and supported by high-confidence fragmented MS/MS spectrum.

### Top cancer peptides presented by HLA-E

We highlighted four HLA-E-presented cancer peptides based on their stability and therapeutic potential (**Fig. 6C,D**). For therapeutic potential, we considered the RNA expression differential between cancer and normal tissues, gene involvement in tumor survival, and the size of the addressable patient population. All four peptides we highlighted have above 50% stability values and above 10 fold enrichment if they originated from our cell based screening assay. Notably, ETV4-KLMDPGSLPPL was discovered by MHC Attention 2.0 in the computational screening benchmark as the 17^th^-highest-ranked HLA-E peptide out of 71,038 across 63 genes (99.99 presentation percentile score).

## Discussion

HLA-E is an attractive target for cancer immunotherapy because of its near-universal allele coverage and its central role in tumor immune evasion through engagement of inhibitory NK-cell receptors (*14*). However, systematic discovery of HLA-E-presented tumor antigens has remained challenging because HLA-E has restricted binding preferences and many predicted non-canonical peptides fail to form stable peptide-MHC complexes. In this study, we addressed this challenge by developing an integrated discovery platform for HLA-E presentable targets that combines mammalian cell-based assays and machine learning. We then used this platform to identify and validate a large number of novel HLA-E peptides, including several that may serve as promising cancer associated targets (**Supplementary Table 2–5**).

Our work builds on prior findings that demonstrate HLA-E presenting peptides can elicit a robust T-cell response (*21, 45*). For example, Lyer et al. reported that vaccinated rhesus macaques can mount strong CD8+ T-cell immunity against human Wilms tumor-1 (WT1) protein in the context of MHC-E (*20*). In this work, we expand on this finding to demonstrate that the most stable HLA-E peptide from WT1 is likely RMFPNAPYL (**Fig. 6C**). Notably, this WT1 peptide is a well-known peptide presented by HLA-A*02 (*46*). Shared peptides between HLA-E and HLA-A*02 have previously been reported (*47*), and may be particularly desirable targets for therapeutic modalities like TCR-T and cancer vaccines.

One striking result from our work is that many reported HLA-E peptides in public databases like IEDB may not stabilize HLA-E complexes on the cell surface. Specifically, more than 63% of IEDB reported HLA-E peptides have an enrichment value lower than 1 in our screening assays. Upon inspection, the large number of these peptides are non-HLA peptides detected in previous MS immunopeptidomics studies (*48, 49*). It is also possible that some of these peptides form stable complexes with HLA-E in intracellular compartments, but are unstable on the cell surface. Current antibody-based MS immunopeptidomics studies don’t differentiate these two populations (*50*), but our cell library based screening assays do.

Our results also highlight the importance of additional HLA-E training data for MHC presentation prediction models. For example, the strongest cancer associated HLA-E presenter (ETV4-KLMDPGSLPPL) was discovered by MHC Attention 2.0 after training on HLA-E library screening data. NetMHCpan4.2 gives this top ETV4 presenter a percentile of 98.78, which is high but far below the peptides it selected (>99.98), likely because the peptide does not follow motifs common to IEDB-reported HLA-E peptides. We see further evidence of this pattern in the length distribution of peptides selected by the models. The peptides selected by NetMHCpan4.2 are all 9mers, whereas the peptides selected by MHC Attention 2.0 HLA-E include many 10 and 11mers (including ETV4-KLMDPGSLPPL). However, the most stable HLA-E presenting 9mer identified (MATR3-ILGPPPPSF) also came from MHC Attention 2.0. NetMHCpan4.2 does give this peptide a high score as well (99.39 percentile), but it is again crowded out by other even higher scoring peptides. This suggests that NetMHCpan4.2 does have useful predictive power for HLA-E, but still gives many false positives at high percentiles (98+). In contrast, likely due to its expanded training data, MHC Attention 2.0 HLA-E appears to have useful discrimination power for true presenters even at very high percentiles, with top 2 precision of 50%, top 5 precision of 40%, and precision over the full top 20 of 25% (**Fig. 6B**).

MHC Attention 2.0 outperforms other MHC presentation prediction models on both muti-allele benchmarks and single-allele benchmarks. This is notable from an architecture standpoint because in the single-allele benchmark there is no allele deconvolution problem for the attention mechanism to solve. One potential explanation is that the MHC encoder learns stronger allele representations during training: in multi-allele examples, the model must distinguish among candidate presenting alleles through the attention mechanism, and the representations learned to support this discrimination may improve prediction even when only a single allele is provided at inference time. More generally, prior work has observed that deep learning models often perform best when trained end-to-end rather than relying on heuristics or feature selection (*51*). In this sense, incorporating multi-allele data directly into model training—as opposed to the allele selection heuristics used by NetMHCpan4.2 and MHCflurry 2.0—can be viewed as an instance of this broader pattern.

The prioritized HLA-E cancer peptides in **Figure 6C** are potentially good candidates for cancer drug development based on our *in vitro* validation studies and broader biological rationale. WT1 is one of the best-established intracellular cancer antigens, with expression across leukemias and multiple solid tumors, as well as longstanding preclinical and clinical support for vaccine-and TCR-based immunotherapy (*52, 53*). Similarly, RNF43 is a clinically meaningful source antigen even though it has not been widely pursued as a pMHC target (*54*). RNF43 is a compelling source antigen because recurrent loss-of-function mutations in gastrointestinal cancers (6-18% of colorectal cancers) can promote Wnt-dependent tumor growth, linking RNF43 alteration to a functional cancer dependency (*55, 56*). BMP8A is less clinically mature but remains notable because prior studies showed its overexpression in thyroid cancer and linked it to tumor progression, poor prognosis, invasion, and drug resistance (*57, 58*). Finally, ETV4 is an oncogenic transcription factor implicated in proliferation, invasion, metastasis, and treatment resistance across multiple tumor types (*59, 60 , 61*). Similar to many transcription factors, ETV4 remains difficult to drug with conventional small molecules or antibody-based modalities. In this context, identifying ETV4-KLMDPGSLPPL as a stable HLA-E peptide, with support from both the peptide stability assay and mass spectrometry, is particularly significant because it converts a biologically compelling but classically hard-to-drug intracellular driver into a tractable pMHC target.

This work has several technical limitations. First, our high-throughput screening assay is designed around peptide-dependent HLA-E stability in an engineered context and does not fully capture endogenous antigen processing and competition. Before adopting HLA-E candidate peptides as therapeutic targets, it is important to evaluate them in competition assays against endogenous VL9 peptides. Second, while mass spectrometry provided additional support for a subset of HLA-E presenting candidates, the overlap between our validated hits and immunopeptidomics was limited. This likely reflects both the technical sensitivity of MS technology and biological differences between stability and endogenous presentation. Finally, it is possible to expand our peptide screening even further using the improved prediction algorithm and improved DNA oligo synthesis. For example, a DNA library size of 50,000 peptides is now tractable using silicon-based DNA microarray synthesis (*62*).

Several areas of future work are suggested by our results. In particular, this study only examined HLA-E presentation and did not address downstream receptor/binder discovery or therapeutic targeting. Future work might determine whether the validated peptides identified here can support productive recognition by TCRs or other binding modalities. Recent advances in protein binder discovery (*30, 63, 64*) and emerging *de novo* protein design (*65–68*) can enable rapid therapeutic development against these HLA-E-presented targets. Beyond HLA-E presentation, this study also did not address whether the peptides we identified interact with NKG2C (activating) or NKG2A (inhibitory) (*14*). For example, peptides that activate NK cells might be preferable for the development of cancer vaccines.

Overall, this study establishes a cancer target discovery platform tailored to the biochemical constraints of HLA-E. The experimental data and screening results presented here both deepen our understanding of HLA-E antigen presentation and also directly facilitate the development of pMHC-targeting therapies. Such therapies are particularly valuable because of the near universal allele coverage of HLA-E, providing a more practical path toward pMHC-targeting studies at global scale (*9, 10*).

## Methods

### Cancer target prioritization and analysis of RNA-seq datasets

We started with genes from the Cancer Antigenic Peptide database (CAPED) (*36*) as these genes have been demonstrated to produce cancer associated MHC peptides. To further supplement the gene list, we analyzed bulk tumor and normal tissue RNA-sequencing datasets from the Cancer Genome Atlas Program (TCGA) (*34*), the Genotype-Tissue Expression (GTEx) (*33*). Gene expression values were harmonized as transcripts per million (TPM) and compared across tumor and corresponding normal tissues. Candidate genes were prioritized based on two predefined criteria. First, genes were required to show at least a fourfold higher expression in tumors (median across TCGA samples) relative to normal tissue controls (median across GTEx samples). Second, genes were required to show expression below 5 TPM in critical normal organs, including brain, heart, lung, kidney, and liver. Genes meeting these criteria were further reviewed manually based on literature support for cancer association, oncogenic function, or prior therapeutic interest.

Single-cell RNA-seq data were obtained from the 3CA database (*35*) and normalized to transcripts per million (TPM) by scaling counts to 1,000,000 per cell followed by log2 transformation with a pseudocount of 1. For each dataset, cells were annotated as malignant (per author annotation) or non-malignant (all other cells). To quantify the ability of each gene to distinguish malignant from non-malignant cells, we calculated the area under the receiver operating characteristic curve (AUC). For each gene, binary class labels were assigned to cells (1 = malignant, 0 = non-malignant), and normalized expression values were used as a continuous predictor. AUC was computed per dataset using the R pROC package (v1.18.5) as a threshold-independent measure of classification performance, with higher expression values corresponding to increased likelihood of malignancy. AUC values were subsequently mean-aggregated across datasets within each cancer type. Only genes with sufficient representation in both groups (≥2 cells per class) were included. A minimal 0.6 AUC value is required for a gene to be considered for further analysis.

Pseudogenes, non-coding genes, and genes with FDA approved drugs were excluded from the analysis. We produced 141 cancer-associated gene candidates combining bulk RNA-Seq analysis, single-cell RNA-Seq, and the CAPED for downstream peptide discovery (**Supplementary Table 1**). A subset of these genes were selected for second round screening with MHC Attention 2.0 HLA-E, focusing on genes of clinical interest or underrepresented in the cell library (n=63).

### Peptide enumeration and candidate sequence generation

Protein sequences corresponding to genes were retrieved from UniProt (*69*). All possible peptides of length 9 (cell-based screening library, **Fig. 3**) or 9-11 (final MHC Attention benchmark, **Fig. 6**) were generated using a sliding-window approach. For library candidate generation, we selected peptides from cancer associated genes with scores above 98th MHC Attention 1.0 percentiles (n = 1051). We also control HLA-E peptides (positive controls, n = 5; negative controls, n=3), IEDB-reported HLA-E peptides (n = 737), predicted peptides from the normal human proteome (n = 3144), and PRAME-derived peptides (n = 445) as an explorative screening. Duplicate peptide sequences were collapsed prior to model scoring. Peptide identifiers were retained throughout the workflow to enable integration of prediction scores, cell-based screening results, and orthogonal validation assays.

### MHC Attention architecture

MHC Attention is a deep neural network for MHC class I presentation that learns jointly from single-allele and multi-allele immunopeptidomics data. Its central design feature is the treatment of the presenting allele in multi-allele samples as a latent variable, with attention weights over the candidate allele set learned during training as a form of pointwise attention. This enables training on large-scale mass spectrometry datasets in which presentation is observed at the sample level, and a set of up to six candidate alleles are known, but the true presenting allele is not labeled.

The model takes a peptide sequence and a set of MHC class I pseudosequences as input. Amino acid tokens are embedded in a shared learned amino acid space, with separate positional embeddings for peptide and MHC inputs. Peptide and MHC sequences are then encoded by independent one-dimensional convolutional encoders with batch normalization and max pooling. The peptide encoder uses convolutional channel widths of 128, 256 and 512, and the MHC encoder uses 64, 64 and 128, and the model takes 6 MHC alleles as inputs to support training on multi-allele data. The encoded peptide representation is concatenated with each candidate allele encoding, passed through a learned allele attention component and normalized with a softmax over candidate alleles. This results in a pooled MHC representation, which is concatenated with the peptide representation and the selector outputs and passed through a final multilayer scoring block to produce a scalar presentation score.

Allele attention here takes the following form, where M_i_ is a candidate among 6 MHC alleles after CNN encoding, P is the target peptide after CNN encoding, and AttentionMLP is a fully-connected feedforward neural network component:

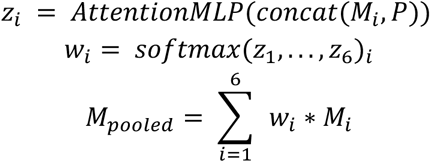

Here z_i_ is the attention score per candidate allele i, w_i_ is the attention weight for candidate allele i, and M_pooled_ is the allele representation used for final scoring after attention weighting.

### MHC Attention training data and preprocessing

Training data were derived from the NetMHCpan 4.1 and 4.2 training releases (*2, 70*) and comprised eluted ligand (EL) and binding affinity (BA) examples. The training split for MHC Attention 1.0 contained 13,075,736 examples across 250 alleles (12,867,643 EL and 208,093 BA), of which 715,519 were positive and 12,360,217 were negative. The training split for MHC Attention 2.0 contained 17,563,641 examples across 273 alleles (17,348,838 EL and 214,803 BA), of which 846,715 were positive and 16,716,926 were negative. Examples contained up to six candidate alleles. Inputs were restricted to peptides of up to 15 amino acids and MHC pseudosequences of length 40.

Allele slots were packed using a repeat-last scheme: when fewer than six alleles were available, the last observed allele was repeated to fill the remaining slots. This scheme was applied consistently during preprocessing, training and deployed inference.

### MHC Attention model training hyperparameters and HLA-E fine-tuning

For both versions 1.0 and 2.0, MHC Attention was trained on all training data using the following hyperparameters: batch size 128, learning rate 1 × 10⁻³, dropout 0.3, 10 epochs, six allele slots, peptide length 15, MHC pseudosequence length 40, embedding dimension 50, selector hidden dimension 128 and combination hidden dimension 256. The BA threshold was set to 0.426, and BA and curated losses were weighted equally (1.0).

To enable HLA-E-specific scoring, we trained a dedicated HLA-E head on top of the MHC Attention 2.0 pre-trained base model. The peptide and MHC encoders from the trained class I model were frozen, and peptide and MHC features were projected into a shared hidden space. The HLA-E head combined peptide features, MHC features, their elementwise product and their absolute difference, followed by a two-layer multilayer perceptron that produced a scalar HLA-E score. Fine-tuning used repeat-last packing, batch size 128, learning rate 1 × 10⁻³, dropout 0.3, and 100 epochs. Targets were derived from normalized HLA-E abundance using a clipped linear transform with a soft minimum of 0.0 and a soft maximum of 50.0, and optimization used weighted binary cross-entropy with logits.

### MHC Attention Prediction Benchmarks

We evaluated models on three validation benchmarks:

#### Multi-allele validation

This benchmark was derived from a filtered subset of the Sarkizova et al. multi-allele immunopeptidomics dataset (*1*). To enable direct comparison with MHCflurry 2.0, we used the overlap-filtered subset shared between the MhcAttention, NetMHCpan and MHCflurry. The benchmark contained 102,728 peptide rows, of which 3,200 were positives (**Supplementary Table 6**).

#### Single-allele validation

This benchmark was constructed from the Netmhcpan 4.1 validation set, assembled from the same source family as the Sarkizova data. This data was again filtered to remove any training overlap between MHC Attention and Netmhcpan. MHC Flurry had nearly complete overlap with this set and so could not be fairly evaluated. This dataset was evaluated per allele and contained 931,487 rows and 31,707 positives across 36 alleles (**Supplementary Table 6**).

#### HLA-E validation

This held-out benchmark was derived from Vcreate’s HLA-E library experiments and contained 1,028 held-out rows (20% of data) used to evaluate all versions of the algorithm, as well as Netmhcpan and MHC Flurry. Positives were defined as peptides with normalized HLA-E enrichment greater than 4, yielding 67 positives (**Supplementary Table 6**).

We compared all versions of MHC Attention with NetMHCpan4.2 and MHC Flurry 2.0 (*27*). MHC Flurry 2.0 was excluded from the single-allele benchmark because of training overlap within this benchmark bundle.

### MHC Attention scores and evaluation metrics

Primary comparisons used raw model scores. For MHC Attention 2.0 and 1.0, this corresponds to the presentation score; for NetMHCpan4.2, the EL score; and for MHCflurry 2.0, the presentation score. All models reported the max score for multi-allele examples. We report area under the receiver operating characteristic curve (AUC), partial AUC truncated at a false-positive rate of 0.1 (AUC0.1) and a top-*k* precision metric denoted PPV. This PPV was computed by ranking predictions within each evaluation slice, taking the top *k* predictions where *k* = 0.95 × (number of positives in the slice) and computing precision within that ranked set. For single-allele validation, we additionally report macro-by-allele summaries, computed by evaluating each metric independently per allele and averaging across alleles.

#### Presentation percentiles

In this work we report presentation percentiles reported by MHC Attention and NetMHCpan4.2 for various peptides. For MHC Attention, these percentiles were calculated by taking the percentile rank of a peptide’s score against a background distribution of 100,000 random human peptides for the query allele. We converted the NetMHCpan4.2 model output “rank” into percentile using the formula: percentile = 100 - rank. Thus both tools report percentiles under the same higher-is-better assumption.

### Selection of candidate peptides for HLA-E screening

Candidate peptides for experimental testing were either HLA-E peptides reported by the IEDB (*32*) or selected based on MHC Attention prediction scores for HLA-E presentation. Peptides derived from the whole human proteome and prioritized cancer-associated genes were ranked separately based on MHC Attention 1.0. The final screening library also included high-scoring candidate peptides together with positive control peptides (n=5), negative control sequences (n=3), and exhaustive 9mer sliding windows of PRAME protein (n=437). PRAME 9mer peptides serve as both negative controls and a focused screening for PRAME, an important cancer testis antigen (*71*). The peptide number in this “6000” peptide library is 5385 as some peptides are introduced in more than one category (e.g. both IEDB reported and predicted). Oligonucleotides encoding the peptide library were reverse translated and codon optimized with DNA Chisel (*72*) and synthesized with the Twist Bioscience Oligo Pools service.

#### Construction of natural HLA-E lentiviral plasmid and HLA-E single-chain trimer lentiviral plasmid

Both natural HLA-E gene and HLA-E single-chain trimer backbones were cloned into a 3rd generation lentiviral backbone plasmid (Twist Bioscience, pTwist Lenti SFFV) with the Twist Bioscience Clonal Gene service. The natural HLA-E gene lentiviral plasmid contains codon-optimized HLA-E gene and human B2M gene linked by a T2A cleavage linker.

HLA-E single-chain trimer constructs were used for peptide stability assay in a mammalian cell-based HLA-E display system (**Fig. 3**). Specifically, each construct encoded signal peptide, test peptide, GGGGS linker, human B2M, GGGGS linker, HLA-E heavy chain (Y84A), and C-terminus mNeonGreen reporter (GFP). Exact linker sequences and domain arrangement were designed based on previous studies (*14*) and provided in the supplementary GenBank file (**Supplementary File 1**). Y84A was introduced to accommodate peptide loading without artificially stabilizing non-presenting HLA-E peptides (reported in Y84C designs). The cloning of HLA-E single-chain trimer lentivirus consists of two steps. The original construct backbone sequence with a peptide cloning site (two PaqCI cutting sites) and all other essential components was first cloned into the Twist Bioscience pTwist Lenti SFFV plasmid with the Twist Bioscience Clonal Gene service. In the second step, peptide DNA oligo sequences were inserted into the constructs using PaqCI-based Golden Gate Assembly according to the manufacturer protocol (NEB, catalog number R0745S).

Plasmid sequences were verified by whole plasmid sequencing (random sampling of E coli plasmids, Genewiz Plasmid-EZ service) and amplicon sequencing (bulk-sequencing, Genewiz Amplicon-EZ service). Peptide frequencies in lentiviral plasmid library were similar to peptide frequencies in the final mammalian cell library.

### Cell culture

K562 cells were obtained from ATCC (catalog number CCL-243) and maintained in RPMI 1640 media supplemented with 10% FBS (Sigma-Aldrich, catalog number F2442), 1% Penicillin and Streptomycin (Sigma-Aldrich, catalog number P4333), and 1% Glutamax (ThermoFisher, catalog number 35050061) at 37 °C in a humidified incubator with 5% CO2. Similarly, HEK293T cells (ATCC, catalog number CRL-3216) were cultured with DMEM medium with the same set of supplements and growing conditions. Cells were routinely tested for mycoplasma contamination using MycoStrip (InvivoGen, catalog number rep-mys-10). For screening experiments, cells were maintained within a density range of 5 x 10^5^ to 10^6^ to minimize variation in transduction efficiency and surface expression.

A HLA-null K562 line was further developed by single-cell sorting parental K562 cells based on their surface HLA-I staining level (APC anti-HLA-I antibodies, BioLegend, catalog number 311410). Individual K562 cell clones were expanded and further profiled for their surface HLA-I level after the clone reached around one million cells. Using a HLA-null K562 cell line minimizes the presence of peptides from HLA-A, -B, -C in the MS immunopeptidomics study.

### Lentiviral production and transduction

Lentiviral particles encoding individual constructs or pooled peptide libraries were produced in HEK293T cells (ATCC, catalog number CRL-3216) by transient transfection with transfer plasmid, packaging plasmid, and envelope plasmid according to the manufacturer protocol (Mirus TransIT Lentivirus System, catalog number 6650). Viral supernatants were collected at day 2 and filtered with 0.45 µm PES syringe filters (NEST Scientific, catalog number 380211). HLA-null K562 cells were transduced at a multiplicity of infection (MOI) of 0.2-0.3 in the presence of 6 µg/mL DEAE-Dextran (Sigma, catalog number 00898). K562 cells transduced with natural HLA-E genes were sorted with the surface HLA-I level and expanded into a homogenous population for downstream mass spec studies (>98% of cells have high levels of HLA-E level). K562 cells transduced with single-chain trimer constructs were directly used for screening assays 7 days post transduction without further selection.

### Flow cytometry-based cell screening and HLA-E surface stability testing

For pooled screening, transduced cells expressing the HLA-E single-chain trimer library were stained with APC-conjugated anti-HLA antibody (clone 3D12, BioLegend, catalog number 311410), together with viability dye DAPI (Sigma-Aldrich, catalog number D9542). Flow cytometry was performed using a SONY SH800 sorter. Sorting gates were defined sequentially on singlets, viable cells, and GFP+ cells (**Fig. 3A**). GFP+ cells were gated/sorted into populations with high HLA-E surface expression and low HLA-E surface expression. GFP negative cells were not sorted as they do not express HLA-E single-chain trimer constructs. Sorted cell populations were collected into DNA/RNA Shield (Zymo Research, catalog R1100-50).

For individual peptide stability testing, peptides were loaded onto peptide-free biotinylated HLA-E monomers (Acro Biosystems, HLM-H82Eg) at a 20:1 molar ratio of peptide to HLA-E monomer. After loading, peptide–MHC complexes were conjugated to streptavidin-coated beads (Micromod, 08-19-403) and subjected to heat stress at 37°C for 60 min. Beads were then stained with an anti-HLA antibody (BioLegend, 311410) for 15 min and washed twice with PBS. Flow cytometry was used to quantify the percentage of HLA-positive beads above the negative-control threshold, defined as the 99th percentile of HLA fluorescence signal from negative-control beads loaded with either CMV-NLVPMVATV or no peptide. This percentage was used as the peptide stability value. Peptides with stability values above 5% were classified as moderately stable, whereas values above 50% were classified as highly stable.

### Sequencing-based deconvolution of pooled screening results

The sorted HLA-E expressing K562 cells were lysed for total RNA extraction with Zymo Quick-RNA Purification Kit (Zymo Research, Catalog number R1054). Peptide specific mRNA was amplified by RT-PCR using SuperScript IV OneStep RT-PCR System (Fisher Scientific Catalog Number 12594025) and primers flanking the peptide-encoding region. Amplicons were sequenced using Illumina MiSeq (catalog number MS-102-2002). Reads were demultiplexed using constant flanking regions, quality-filtered, and matched to the reference peptide library. Peptide abundance in each sorted population was quantified as normalized read counts. Enrichment score for each peptide was calculated as follows,

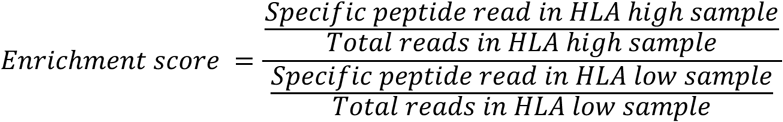

### Immunoprecipitation and mass spectrometry-based HLA-E peptide profiling

For immunopeptidomic analysis, HLA-null K562 cells were transduced and sorted to express HLA-E (see described in the earlier **Method** section). Ten to thirty million HLA-E K562 cells were harvested, washed with PBS, pelleted, dry frozen, and stored at −80 °C until processing. HLA peptide isolation and mass spectrometry analysis were performed by Alithea Bio (Freiburg, Germany) using their established HLA immunopeptidomics workflows (*73*). Briefly, frozen cell pellets were lysed under non-denaturing conditions, and HLA class I complexes were enriched from clarified lysates by immunoaffinity purification using a W6/32 pan-HLA class I antibody coupled to resin. Bound peptide–HLA complexes were washed and acid-eluted, and peptide-containing fractions were separated from the HLA heavy chain, β2-microglobulin, antibody, and other higher-molecular-weight proteins before desalting and concentration for LC–MS/MS analysis. W6/32 antibodies capture both classical and non-classical HLA class I complexes, the resulting peptide set was interpreted as an HLA class I-enriched immunopeptidome from HLA-E-expressing K562 cells rather than an exclusively HLA-E-specific pull-down. Since the original HLA-null K562 cells have minimal HLA expression, most of HLA peptides are expected to be from the high level of transgenic HLA-E complexes.

Purified peptide fractions were analyzed by liquid chromatography–tandem mass spectrometry (LC-MS/MS) on a timsTOF Ultra mass spectrometer in data-dependent acquisition mode. MS/MS spectra were searched against the human UniProt reference proteome using database search software with no-enzyme specificity, consistent with the endogenous non-tryptic nature of HLA-bound peptides. Peptide-spectrum matches were filtered using false-discovery-rate control (1%), and candidate HLA-associated peptides were evaluated based on peptide length, sequence features, and consistency with expected HLA class I immunopeptidome characteristics. 8-12AA long peptides recovered from MS were clustered using GibbsCluster (*44*) to identify peptide sequence motifs. Peptides more than 13 amino acids in length were excluded from the final analysis as they were likely cell lysis contaminants (**Supplementary Table 3**).

#### Statistical test and plotting

Unless specified, statistical significance in this study was determined with two-tailed two-sample unequal variance t-test. Significance of prediction model performance difference across single MHC allele data was determined with paired Wilcoxon signed-rank tests. Plots were generated with python matplotlib. Schematic illustrations were generated using BioRender or HTML.

## Code and data availability

Detailed experimental results are listed in the **Supplementary Table 1-7**. MHC class I training and validation data is available via the original NetMHCpan paper and website (*2*). HLA-E screening data for training and validation is in the **Supplementary Table 2**. The exact sequence of HLA-E single-chain trimer backbone plasmid is provided in the GenBank file (**Supplementary File 1**). MHC Attention 2.0 can be accessed online via https://vcreate.io/mhcattention.

## Author Contributions

B.C., E.F., and M.D. conceived the study, designed the overall research strategy, executed the experiments, analyzed the results, and wrote the manuscript. G.S.G. performed gene prioritization and data analysis. M.K. contributed to experimental execution. B.C. supervised the work. All authors reviewed and approved the manuscript.

## Acknowledgements

This work was partially supported by NSF SBIR grant (2213253, 2406662), NIH SBIR grant (NIGMS, R43-GM143955), and NCI Concept Award (75N91024C00001). G.S.G. is supported by NIH grant T32CA009172. We thank Karan Raj Kathuria for discussions about the manuscript.

## Supplementary Table Index

1. Top cancer associated protein-coding genes
2. HLA-E antigen library with design, sequence and enrichment
3. MS output from HLA-E K562 cell immunopeptidomics
4. HLA-E peptide stability results for Figure 5A
5. HLA-E peptide stability results for Figure 6B
6. Validation Data for MHC Attention statistics
7. Summary table of performance in three benchmarks especially single-allele (Figure 4B)

